# Lyophilized cell-free systems display tolerance to organic solvent exposure

**DOI:** 10.1101/2020.06.11.121418

**Authors:** Marilyn S. Lee, Rebecca M. Raig, Maneesh K. Gupta, Matthew W. Lux

## Abstract

Cell-free systems offer a powerful way to deliver biochemical activity to the field without cold chain storage. These systems are capable of sensing as well as biosynthesis of useful molecules at the point of need. So far, cell-free protein synthesis (CFPS) reactions have been studied as aqueous solutions in test tubes or absorbed into paper or cloth. Embedding biological functionality into broadly-used materials, such as plastic polymers, represents an attractive goal. Unfortunately, this goal has for the most part remained out of reach, presumably due to the fragility of biological systems outside of aqueous environments. Here, we describe a surprising and useful feature of lyophilized cell-free lysate systems: tolerance to a variety of organic solvents. Screens of individual CFPS reagents and different CFPS methods reveal that solvent tolerance varies by CFPS reagent composition. Tolerance to suspension in organic solvents may facilitate the use of polymers to deliver dry cell-free reactions in the form of coatings or fibers, or allow dosing of analytes or substrates dissolved in non-aqueous solvents, among other processing possibilities.

## Introduction

Cell-free systems are a collection of techniques for activating cellular machinery outside of living cells. Key cell-like functionalities in these systems include cell-free protein synthesis (CFPS)(1–4) and complex metabolism that can both provide energy to the system and produce small molecules of interest (5–9). In these reactions there is no need to maintain cell growth or viability, and removal of the cell membrane facilitates direct addition or measurement of molecules like DNA. These systems have been used for a wide range of applications, including sensing, manufacturing, genetic prototyping, and education (10). There are two main approaches to reconstitute cellular activity *in vitro* (Figure 1A). One approach is the use of crude cell lysates, where cells are broken open and additional building blocks and buffers are added to allow the proteins and other components in the lysate to retain much of their native function (3, 11). Since many undefined components from the cell are still present, there is limited control over biochemical activity. Another approach is the PURE system, where the protein components essential for transcription, translation, and energy cycling are purified before the mixture is reconstituted with energy and substrate solutions (12). This system is much better defined, though it is more expensive to implement and lacks any metabolic enzymes present in lysate that are not individually identified and purified to be included in the reaction.

**Figure 1.**
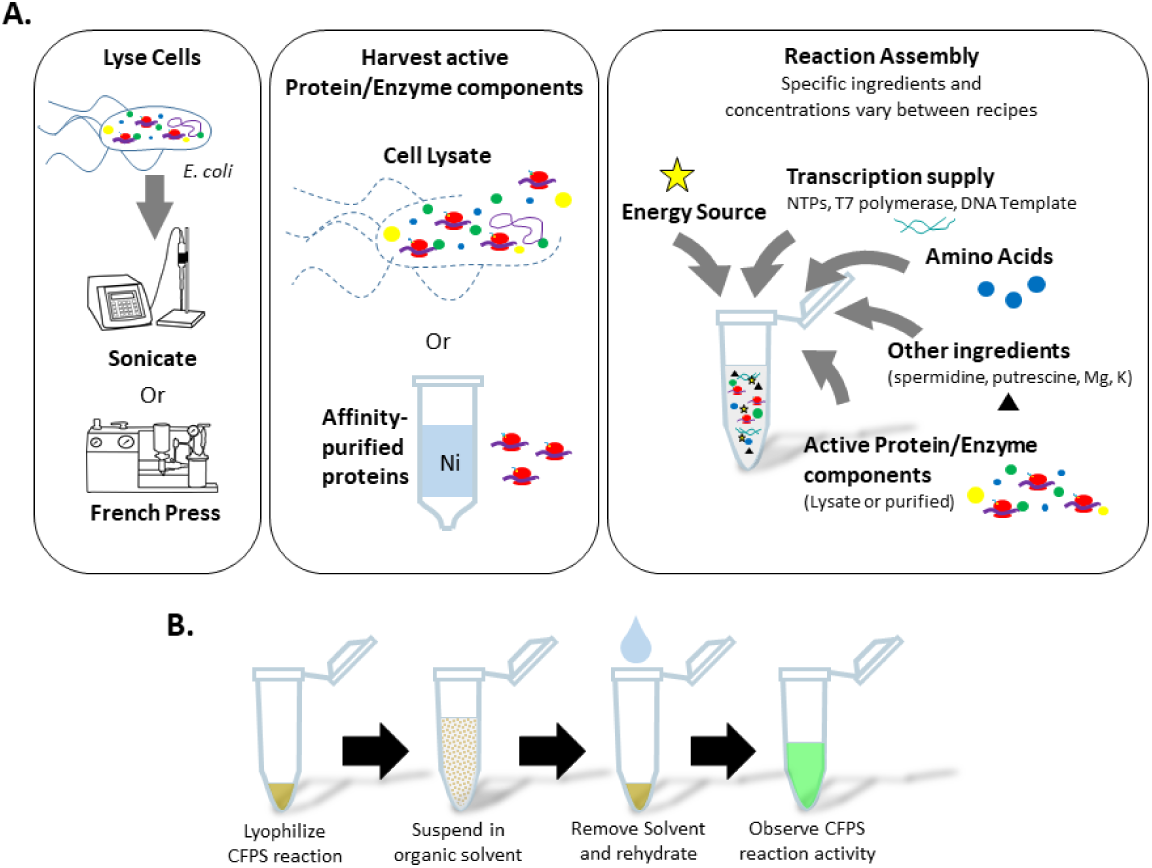
(A) Illustration of the composition of a CFPS reaction and variation in published protocols among lysis methods, purity of active enzyme components, and buffer ingredients. (B) Treatment sequence for CFPS reactions includes lyophilization, suspension in an organic solvent, removal of the organic solvent, rehydration, and assessment of CFPS activity via monitoring the appearance of GFP.

A major emerging application of CFPS reactions is rapid field-deployed sensing (13–17) or molecule production at the point of need (17, 18) with minimal required equipment. Though many biological reagents are normally stored frozen or refrigerated, CFPS reactions may be lyophilized for room temperature storage and retain high levels of protein synthesis activity upon rehydration (19–21). CFPS may be used to quickly prototype and implement gene circuit designs that can sense analytes via a variety of mechanisms and produce human readable outputs such as a colorimetric, fluorescent, or even electrochemical reporters (22). Drying CFPS reactions onto paper tickets or cloth has greatly broadened the scope of applications for cell-free techniques (13, 14, 16, 23, 24). Paper-based reactions retain activity, enabling formats similar to pH paper that are less cumbersome for field use. One draw-back to CFPS reactions is that short reaction lifetimes of a few hours make these systems one-use only. A second draw-back is the need to manually add water to activate the lyophilized reaction. Turning to material science to once again redesign the reaction environment could alleviate these issues.

There is great potential to leverage materials to modulate hydration, and therefore control activation of cell-free systems. Polymers have been widely used to control release timing for cargos that include anything from pharmaceuticals to fertilizers (25, 26). The same principle may be used to extend the lifetime of activity of a detection device. For instance, continual activation of fresh bio-active cargo, such as enzymes, embedded in a material may be achieved by polymer erosion (27, 28). However, this type of formulation requires the biological molecule to be cast into polymers that have low solubility in water, likely necessitating the application of heat or organic solvents. These treatments have a deleterious effect on the activity of most enzymes unless their structure can be protected in some way (29). For a complex mixture like CFPS reactions, it would be reasonable to suspect that such effects on any of the numerous enzymes and other constituent components could result in reduction or elimination of overall protein production.

Non-aqueous enzymology is the study of protein behavior in non-aqueous organic solvents. Over the past few decades, researchers in this field have shown that a subset of purified proteins can, under certain circumstances, maintain their fold and even activity when suspended in organic solvents (30, 31). An important factor influencing protein behavior in organic solvents is the amount of water present in the mixture. Too much water will allow the protein to change conformations, leading to irreversible folding and inactivation. This has been seen in studies titrating organic solvents into aqueous bio-mixtures such as a CFPS reaction (32, 33). Therefore, freeze-drying or spray-drying is necessary to carefully remove water from an enzyme sample before solvent addition to better maintain enzyme activity. Indeed, a crystal structure obtained for subtilisin in pure acetonitrile demonstrates that protein structure in this solvent is very similar to the native aqueous structure (31). Tolerance to non-aqueous solvents has been studied for a small subset of purified proteins, but has never before been explored for a complex bio-mixture like a CFPS reaction in which many proteins, nucleic acids, and metabolites must be preserved to maintain transcription and translation activity.

In this work we describe the effects of exposing lyophilized CFPS reactions to organic solvents. We find that CFPS activity is recovered upon rehydration after exposure of the dried reaction to a variety of organic solvents, though some polar solvents such as ethanol reduce activity. Screens of CFPS reaction recipes, reagent mix components, and solvent removal methods indicate that tolerance to certain solvents is dependent on the presence of additives. Further, we discuss how solvent tolerance of CFPS systems may open the door to new applications by embedding dried CFPS components in new types of polymer materials via solvent casting or sensing analytes in non-aqueous samples.

## Results and Discussion

### Solvent screen

Initial experiments exposing CFPS reactions to organic solvents utilized an *E. coli* lysate system closely mimicking the recipe constructed by Jewett *et al*. (11) and referred to as PANOx-SP. A panel of solvents including acetone, acetonitrile, chloroform, dichloromethane (DCM), dimethylformamide (DMF), dimethylsulfoxide (DMSO), ethanol, ethyl acetate, methanol, and tetrahydrofuran (THF) were used to challenge both the *E. coli* lysate component lyophilized alone, and the complete lyophilized CFPS reaction mixture. The treatment sequence is summarized in Figure 1B. Solvent properties are summarized in Supplementary Table S1. Each solvent was mixed with the dry cake in microplate wells to form a suspension, then incubated for 1 hour at room temperature. The dried CFPS cake material is largely insoluble in all the organic solvents tested, with insoluble particulates settling quickly after mixing. The solvents were removed by aspiration after brief centrifugation to settle the CFPS material into a pellet. Residual solvent was removed by evaporation at reduced pressure. Then, the reactions were rehydrated with water to reach the desired reaction volume. For samples in which only the lysate is dried and exposed to solvent, the other components of the CFPS reaction were added in the form of aqueous solutions at the rehydration step to initiate protein synthesis activity.

CFPS activity was assessed by tracking GFP production from a plasmid DNA template with a T7 promoter (Figure 2, Supplementary Figure S1). We note here that variability of some experimental replicates in this study are higher than in other recent CFPS works that report 6.8-11% variability for individually mixed replicates (34–36). We attribute this difference to either variable loss of some particulate during solvent aspiration or variable penetration of the solvent into particulates during exposures. GFP productivity after organic solvent exposure clearly has some dependence on both the properties of the solvent and the components included during drying and solvent exposure. When the complete reaction is exposed to organic solvents, acetonitrile and ethyl acetate exposure did not result in any loss of activity; acetone, THF, Chloroform, and DCM caused partial loss in activity; and ethanol, DMF, DMSO, and methanol caused total or near total loss of activity. When lysate alone is exposed to solvents, a different pattern emerges: acetonitrile and ethyl acetate again show no loss of activity; acetone, ethanol, chloroform, DMF, methanol, and DCM all yield varying levels of activity; and DMSO and THF completely deactivate the un-supplemented lysate. It is notable that, depending on the solvent type, activity may only be lost for the complete reaction (DMF, Methanol, Ethanol) or only for the lysate (THF), suggesting multiple mechanisms for CFPS inactivation are present.

**Figure 2.**
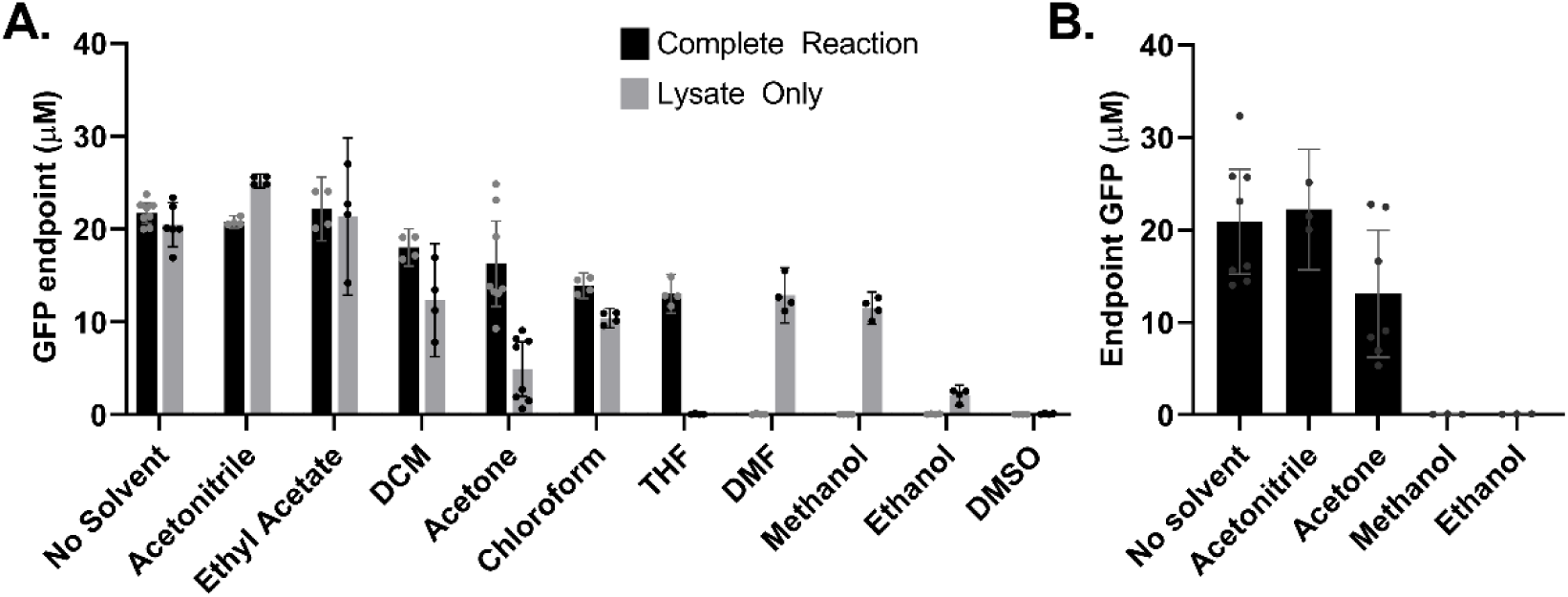
(A) Endpoint concentration of GFP in reaction mixtures after lyophilized lysate (grey bars) or complete CFPS reactions (black bars) were exposed to an organic solvent and rehydrated. In the “no solvent” control the dried reactions were rehydrated without solvent exposure. Error bars represent the 95% confidence interval. (B) Activity post-solvent exposure and evaporation at ambient conditions.

There are several possible ways treatment with an organic solvent might impact the productivity of a CFPS reaction. First, the solvent could cause protein unfolding in the lysate or supplemented polymerase and RNase inhibitor protein components in the dry state. Second, the solvent could extract critical resource molecules from the dried cake into solution and cause them to be removed from the system when the solvent is aspirated away. Third, residual solvent not sufficiently removed by evaporation could cause protein unfolding or other inhibition when water is re-introduced (32). Depending on the properties of each solvent, a different combination of these effects may have an impact.

We investigated whether specific solvent properties correlated with CFPS productivity results (Supplementary Figures S2 and S3, and Supplementary Tables S2 and S3). Statistically significant correlations were observed for complete reaction exposures but not for lysate alone. These correlations could prove informative for selection of additional solvents. For example, low solvent hydrogen bonding propensity correlated well with higher productivity. Still, due to the complexity of potential solvent interactions with the components of the CFPS reaction, prediction of the compatibility of different solvents or solvent blends is difficult without experiment. Thorough discussion of correlation results may be found in the supplementary information.

Further experiments were designed to investigate the impact of solvent removal method and lyophilization at larger scale. For some applications, it may not be possible to apply a vacuum to evaporate solvent at reduced pressure. We found that application of a vacuum was unnecessary for maintenance of complete CFPS reaction activity after exposure to acetonitrile or acetone (Figure 2B). Further, washing with volatile acetonitrile did not improve activity after exposure to less-volatile DMSO (Supplementary Figure S4). These experiments indicated that solvent tolerance was not very sensitive to solvent removal method, at least for the subset tested. Applications such as casting dried CFPS into a material may require much larger quantities of lyophilized CFPS powder than the microplate scale used for screening. We lyophilized larger (250 µL vs 15 µL) batches of PANOx-SP CFPS reactions and exposed them to acetone. We found that CFPS activity was maintained (Supplementary Figure S5). Thermogravimetric analysis (TGA) was a useful technique to confirm a water content of less than 3% (wt/wt). Further discussion of these results may be found in the supplementary information.

### CFPS reaction component screen for acetone tolerance

The observation that the complete dried CFPS reaction performed better than the lysate alone after exposure to acetone, chloroform, or THF suggested that some component of the resource mix might act as a protective additive. To test this hypothesis, components of the resource mix were combined with lysate individually and screened for protective effects upon exposure to acetone and removal of the solvent without a vacuum. In this experiment, three “components” are themselves aqueous mixtures. “10xSS” contains magnesium, potassium, and ammonium glutamate. “15xMM” contains ATP, GTP, UTP, CTP, folinic acid, and tRNA. “20AA” is a mixture of all 20 canonical amino acids. For experimental expediency, samples were rehydrated with a reagent mix containing all reagents, resulting in doubled final concentrations for the individual component screened. Interestingly, only the 15xMM and PEP resulted in statistically significant difference in the productivity due to increased concentration without acetone exposure (Figure S6). Activity decreased in both cases, either because of the change in final concentration or because the component has some negative effect on the lysate during the lyophilization process. After acetone treatment, CFPS ingredients each have varying relative effects on GFP productivity (Figure 3). Plasmid DNA and HEPES buffer both significantly stabilize the lysate. Remarkably, the 10xSS component alone is sufficient to stabilize the lysate to acetone exposure as well as the complete reaction mixture. This experiment is the first confirmation that altering the composition of the lyophilized CFPS reaction mixture greatly impacts tolerance to organic solvent exposure.

**Figure 3.**
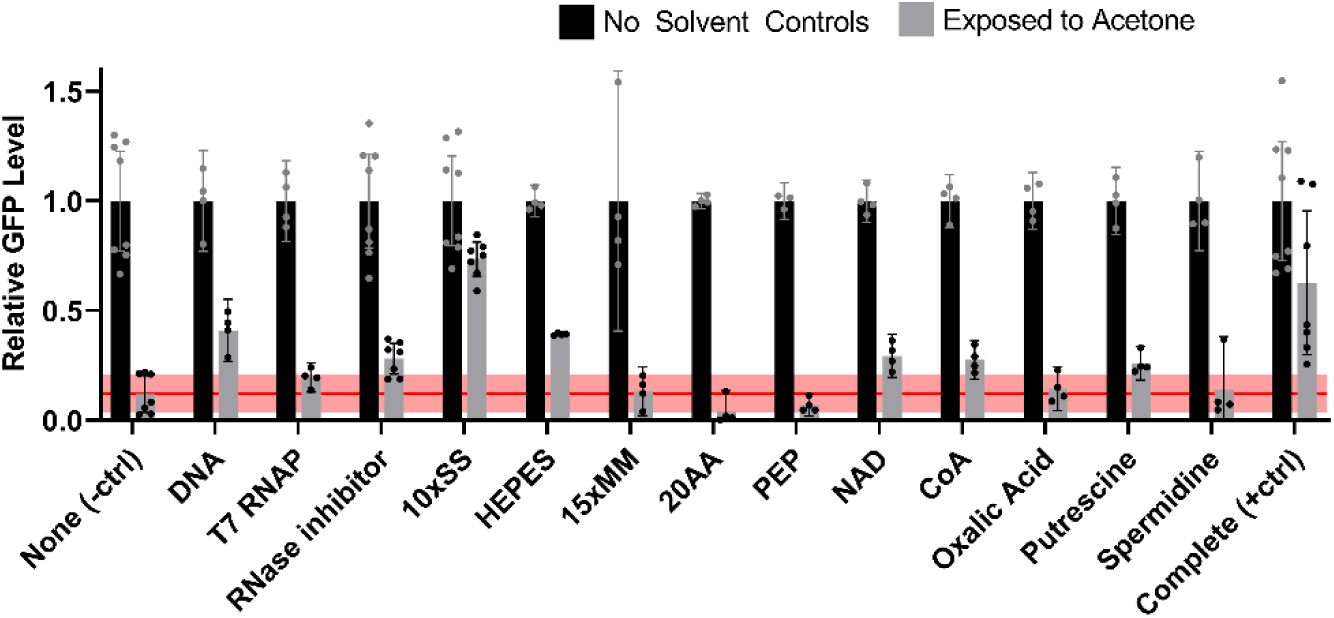
Screen of CFPS components for protective effect during acetone exposure. Black bars are results from controls without solvent exposure. Grey bars refer to a 1 hr acetone exposure. Acetone is evaporated under ambient conditions in this experiment. Each set of bars is labeled with the CFPS ingredient used to supplement the lysate sample during drying. The negative control labeled “None” is lysate without any supplemented additive. The positive control labeled “Complete” includes all CFPS ingredients. The red reference line coincides with the mean fraction of GFP productivity of an un-supplemented lysate control exposed to acetone, with lighter red shading representing the 95% confidence interval. GFP levels are normalized to the no solvent control treatment for each component. Figure S6 depicts the data without normalization.

### Comparing the solvent tolerance of different published CFPS styles

The clear dependence of solvent tolerance on the composition of the reaction led us to investigate solvent tolerance across different published CFPS systems. We tested the performance of three systems: the commercial PURExpress system based on the PURE approach described above (12), the PANOx-SP system used in all experiments above (11), and the 3-PGA system which uses a different strain of *E. coli*, lysis method, and recipe for reaction additives (3, 37). Supplementary Table S4 is a side-by-side comparison of the composition of the CFPS reaction for each recipe. In each case, the reactions are lyophilized and exposed to solvent as described above, with four solvent conditions tested: acetone, acetonitrile, chloroform, or no solvent.

The GFP productivity results confirm that alternate CFPS recipes respond to solvents in different ways (Figure 4, Supplementary Figure S7). Compared to the lysate-based CFPS reactions, the PURExpress formulation has less starting productivity but also minimal sensitivity to treatments with the three solvents tested. Chloroform was the only solvent that significantly reduced GFP productivity in the PURExpress system. On the other hand, the two lysate-based CFPS recipes have very different susceptibility to the organic solvents. 3-PGA is much more susceptible to inactivation by solvent exposure across all three solvent types compared to PANOx-SP.

**Figure 4.**
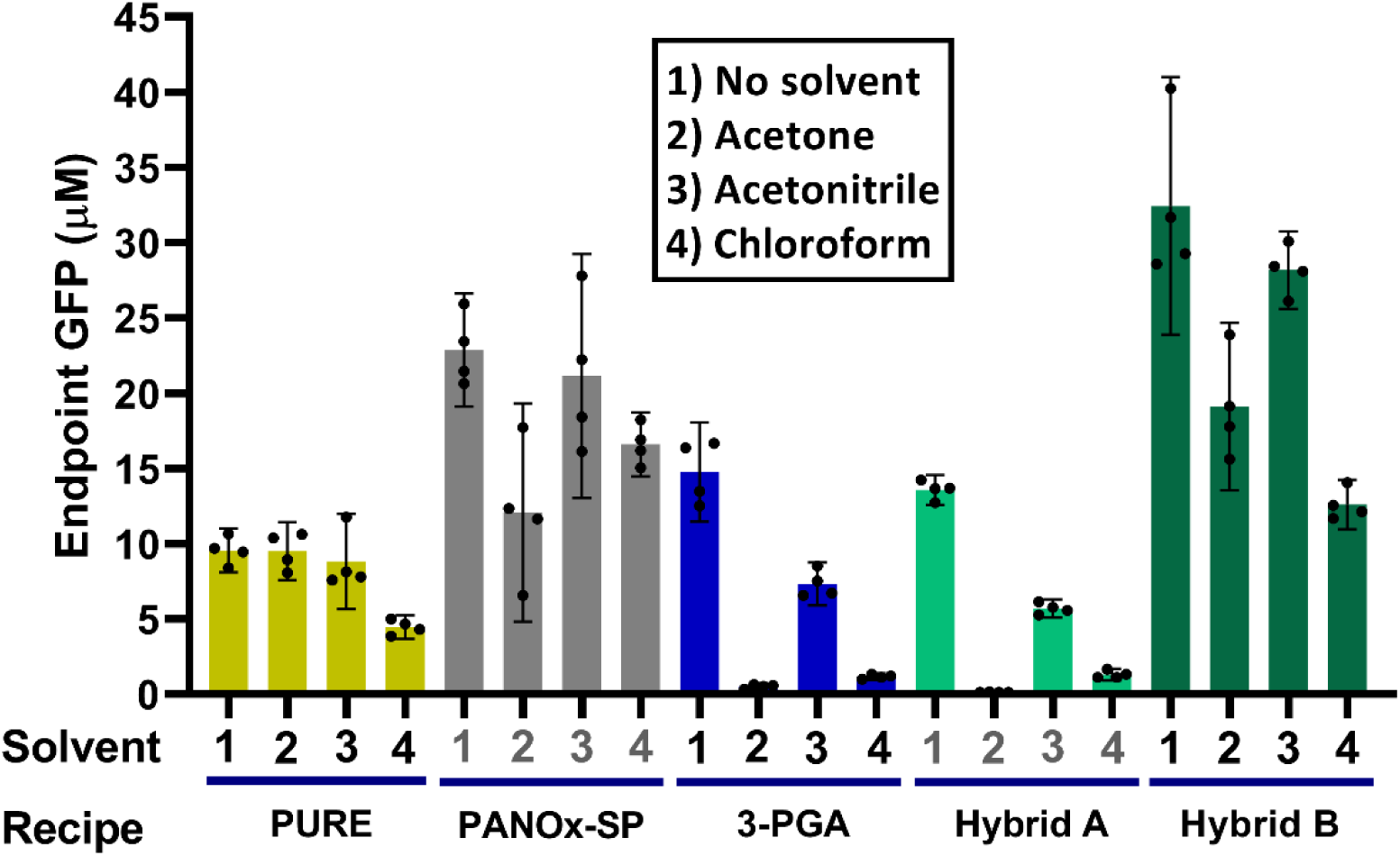
Comparison of different CFPS recipes challenged with organic solvents. Endpoint GFP concentrations are compared for five CFPS recipes after exposure to organic solvents. Error bars represent the 95% confidence interval. Recipe labeled “Hybrid A” is a CFPS reaction mixing PANOx-SP *E. coli* extract and 3-PGA reagent mix. “Hybrid B” is a mixture of 3-PGA extract and PANOx-SP reagent mix.

To learn what components of the CFPS reaction are responsible for increased solvent sensitivity, the *E. coli* extract and reagent mix components of PANOx-SP and 3-PGA CFPS recipes were swapped so that PANOx-SP extract and 3-PGA reagent mix were combined in the “Hybrid A” mixture and 3-PGA extract and PANOx-SP reagent mix were combined in the “Hybrid B” mixture. These hybrid CFPS recipes were then lyophilized, treated with solvent, and rehydrated as before. The results of this test clearly show that samples containing 3-PGA reagents are more susceptible to solvent inactivation than samples containing PANOx-SP reagents. When combined with PANOx-SP reagent mix, 3-PGA extract performed very similarly to PANOx-SP, indicating that differences in the cell extract were not the primary factor contributing to solvent sensitivity. While our focus here was on differences in solvent tolerance and not relative productivity of each system, the hybrid mixtures offer an interesting opportunity to interpret productivity trends in the no solvent cases. It is first worth noting that our yields here are lower than those in the literature for each case, which could be a result of the lyophilization step, unintentional differences in protocol execution, DNA construct, reaction conditions, etc. Nonetheless, we observed that despite no significant difference for the base PANOx-SP and 3-PGA systems, we saw a significant decrease from PANOx-SP to Hybrid A, and a significant increase from 3-PGA to Hybrid B. This result indicates that, at least for our specific implementation and conditions, the PANOx-SP reagent mix yields more protein than 3-PGA reagent mix. Among many differences in the supplemented reagents, a major difference is the type of energy source. PANOx-SP makes used of phosphoenolpyruvate (PEP) to regenerate ATP in the reaction, while 3-PGA utilizes maltodextrin to energize the reaction through substrate phosphorylation and glycolysis. Further experimentation is needed to elucidate specific components leading to solvent sensitivity.

### Summary

The study of the interaction between cell-free reaction components and organic solvents revealed several new findings. Lyophilized CFPS reactions tolerate exposure to a variety of organic solvents without loss or with only partial loss of transcription and translation activity. The degree to which activity is lost by solvent exposure depends on the type of solvent and the ingredients in the lyophilized mixture. Experiments comparing different CFPS recipes indicate that differences in the cell extract preparation are less important for solvent tolerance than the composition of the other ingredients in a CFPS reaction. For acetone exposure, 10xSS alone was sufficient to provide solvent tolerance to the *E. coli* lysate, and DNA and HEPES buffer components provided partial protection.

These newly-discovered characteristics of cell-free systems open the door to many previously elusive applications and directions of study. For instance, the ability to process the biological components of cell-free systems in organic solvents without loss of activity is likely to enable casting these systems into polymeric matrices such as polyurethane that do not dissolve in water. It may also be possible to dose a cell-free sensor system with non-aqueous substrates or analytes. Further study of cell-free systems’ interactions with organic solvent could lead to fundamental insights in non-aqueous enzymology and protein folding. The tolerance of CFPS to solvent when lyophilized raises the question of whether complex systems like the ribosome can maintain activity in an organic solvent either naturally or through engineering. The ability to study these processes in a non-native context is an advantage unique to cell-free systems.

## Materials and Methods

### Reagents

The vendor and catalog number for each organic solvent used in this study is provided in Supplementary Table S1 describing solvent properties. Unless otherwise noted, all other reagents were purchased from Millipore Sigma, St. Louis, MO. Commercial PURExpress kits were purchased from New England Biolabs, Ipswich MA. The plasmid template for expression of GFP via CFPS is PY71sfGFP with genbank accession number MT346027 (38). Plasmid DNA was purified from transformed *E. coli* using a Promega PureYield plasmid midiprep kit, followed by ethanol precipitation to further concentrate and purify the DNA. DNA is stored in RNase and DNase-free water at −20°C until reaction assembly.

### CFPS reaction preparation

PANOx-SP cell extract is prepared from shake flask cultures of *E. coli* BL21 Star (DE3) according to the growth and sonication protocol detailed previously by Kwon and Jewett (11). 3-PGA cell extract is prepared from shake flask cultures of *E. coli* BL21 Rosetta2 and lysed by pressure with a Microfluidizer cell homogenizer. The growth and lysis methods were based closely on protocols published previously by Noireaux et al (3, 37).

Lysates were aliquoted, flash frozen in liquid nitrogen, and stored at −80°C until use in CFPS reaction assembly. To assemble reactions, CFPS components including PURExpress kit solutions, cell lysates, and reagent stock solutions, were thawed on ice, then combined with DNAse free water to reach the final concentrations listed in Supplementary Table S4. Reaction mixtures were well mixed and distributed 15 µL per well into 96 well, v-bottom, polypropylene plates (Costar 3357). Further details on the preparation of CFPS lysates and reagent mixes for PANOx-SP and 3-PGA recipes, as well as lyophilization methods are described in the supplemental information.

### Solvent treatment and rehydration of CFPS reactions

All solvent treatments are performed in a chemical fume hood. 100 µL of each solvent was added to the lyophilized CFPS reaction mixture in designated wells. No solvent was added to lyophilized control reactions. Wells were sealed with a flexible polypropylene mat (Costar 3080) to prevent evaporation. Reactions were incubated with solvent for one hour at room temperature. Following solvent incubation, plates were briefly spun at low speed in a centrifuge to settle the insoluble CFPS reaction components, then solvents were removed by aspiration with a pipette without disturbing the pellet. The residual solvent was removed by evaporation. This was achieved either by allowing the plate to sit uncovered in the fume hood for ambient evaporation, or applying a vacuum at room temperature using a vacuum oven with the heating element turned off for 20 minutes. After solvent removal, all reactions were rehydrated with 15 µL of DNase free water to return CFPS reaction components to their original aqueous concentration. Reaction assembly and solvent treatment methods for the PANOx-SP CFPS component screen are described in the supplemental information.

### Monitoring GFP formation via microplate reader

Immediately after rehydration, plates were sealed with a polypropylene mat, transferred to a BioTek Synergy H1 microplate reader, and incubated at 30°C for 8 hours. Formation of GFP fluorescence e was monitored with ex/em: 485/528 nm with a gain of 100. GFP readings in RFU were converted to µM GFP using fluorescence measurements of purified GFP standards.

## Abbreviations

CFPS: cell-free protein synthesis
GFP: green fluorescent protein
DCM: dichloromethane
DMF: dimethylformamide
DMSO: dimethylsulfoxide
THF: tetrahydrofuran

## Acknowledgements

We thank Stephanie Cole for providing materials for the 3-PGA experiments. We also thank Michael Jewett’s laboratory at Northwestern University and Vincent Noireaux’s laboratory at University of Minnesota for sharing advice and detailed protocols. This work was made possible by funding from the Office of the Secretary of Defense’s Applied Research for the Advancement of Science and Technology Priorities program. Follow-on funding was provided by the CCDC CBC Surface Science Initiative. This work was completed while author Marilyn Lee held an NRC research associateship supported by the CCDC CBC Biological Engineering for Applied Materials Solutions (BEAMS) program.

## Supporting Information

Additional methods details; kinetics data; data and additional interpretation for solvent property correlations, removal of residual solvent by washing, and increased lyophilization volumes

## For Table of Contents Only

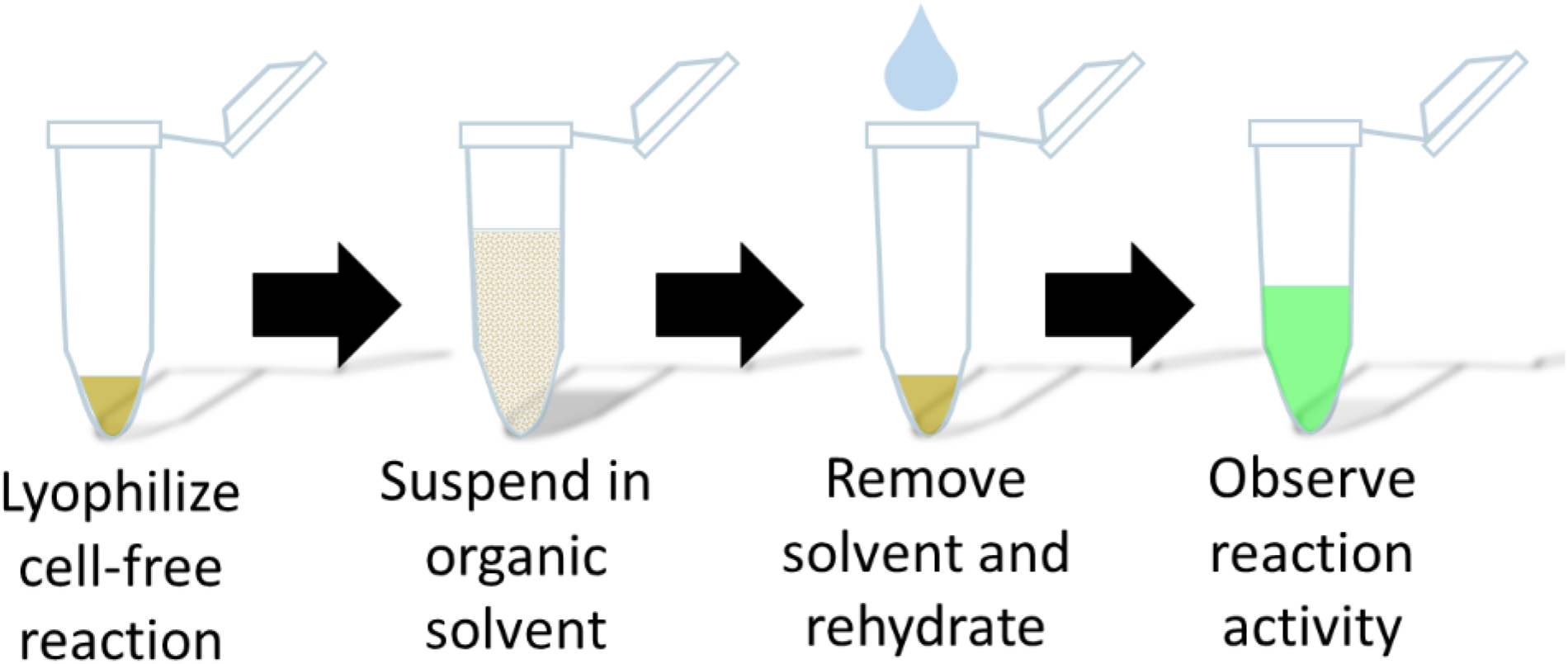

