## Supplementary Information for "Lyophilized cell-free systems display tolerance to organic solvent exposure"

Supplemental Information

Marilyn S. Lee[a], Rebecca Raig[b,c], Maneesh K. Gupta[b], Matthew W. Lux[a*]

[a]US Army Combat Capabilities Development Command Chemical and Biological Center, 8567 Ricketts Point Road, Aberdeen Proving Ground, MD 21010 USA, [b]US Air Force Research Laboratory, 2179 12th St., B652/R122 Wright-Patterson Air Force Base, OH 45433 USA, [c]UES Inc., 4401 Dayton-Xenia Rd., Dayton, OH 45432 USA

[*]Corresponding author:

Matthew W. Lux

Correspondence: 8567 Ricketts Point Road, Bldg E3549 Room B140, APG, MD USA 21010

Supplementary Information

### Supplementary Results and Discussion

*Lyophilizing larger batches of CFPS reaction mixture and measuring water content*

Larger 250 µL scale batches of PANOx-SP CFPS reaction were lyophilized for 48 hours to achieve sufficient dryness to allow milling into a powder (Supplementary Figure S5A). Lyophilization for only 20 hours resulted in a viscid cake that could not be milled and was difficult handle, though still demonstrated normal CFPS activity when rehydrated (data not shown). After 48-hr drying, the dry weight of the powder stabilized at approximately 76 mg per 1 mL of aqueous CFPS solution. Thermogravimetric analysis (TGA) is another more sensitive technique to quantify the amount of water remaining in the sample and only requires around 5 mg of sample for the measurement. TGA measurements indicate that 250 µL samples lyophilized for 48 hours were between 1.2% and 3% water by weight (Supplementary Figure S5B). Endpoint GFP productivity was the same after lyophilization and exposure to acetone compared to a freshly prepared CFPS reaction (Supplementary Figure S5C).

*Correlation of Solvent Properties and CFPS productivity*

Correlation analysis comparing GFP productivity to each of several parameters describing the physical and chemical properties of solvents is summarized in Supplementary Figures S2 and S3, and Supplementary Tables S2 and S3. This analysis sheds light on the trends in solvent characteristics that affect CFPS activity. Supplementary Figure S2 depicts correlation plots for samples where complete CFPS reactions were exposed to solvents. Significant Pearson correlation coefficients were found comparing productivity and solvent boiling point, dielectric constant, Hansen polar parameter, Hansen hydrogen bond parameter, and the octanol/water partition coefficient (log Kow). However, when lysate alone was exposed to organic solvents (Supplementary Figure S3), there were no clear trends between productivity and solvent properties except for a slight but significant relationship to the Hansen dispersion parameter. For complete reaction exposures, decreased solvent hydrogen bonding propensity or increase hydrophobicity correlates with improved productivity, suggesting solvents that participate in molecular interactions similar to water could cause a greater extent of deactivation likely by an increased ability to strip water molecules crucial to protein function. Some polar solvents like ethanol also have a greater tendency to absorb water from the air, which could introduce sufficient concentrations of water to allow protein unfolding or hydrolysis of molecules in the resource mix. There appears to be a cutoff value for solvent boiling point. Two solvents with a high boiling point, DMF and DMSO, caused inhibition of GFP productivity. This could be an indication that trace amounts of these less-volatile solvents are not sufficiently removed by evaporation.

*Solvent evaporation at ambient conditions*

A reduced solvent screen was also implemented without applying a vacuum to learn how evaporation at ambient conditions could affect solvent compatibility. Acetonitrile, acetone, methanol and ethanol were tested on complete lyophilized CFPS reactions. The results are depicted in Figure 2B. GFP productivity is not significantly different for the conditions tested as compared to samples dried with a vacuum.

*Acetonitrile washes to remove DMSO do not improve exposure outcomes*

Since DMSO has a relatively high boiling point of 189°C, it is not clear that evaporation with or without application of a vacuum would be sufficient to remove the solvent from the system in the time allotted before rehydration. It is possible that DMSO causes inactivation of the reaction by protein unfolding or resource extraction, or inhibition is caused by residual DMSO left behind during the rehydration step. To ascertain the real cause, the removal of DMSO was improved by washing with a more volatile solvent. In this test, lyophilized CFPS components were suspended in DMSO, and most of the DMSO was removed by aspiration as before. Then, acetonitrile was used to wash the insoluble CFPS components twice, followed by removal of the acetonitrile by aspiration and application of a vacuum. The activity of GFP expression upon rehydration following this treatment is depicted in Supplementary Figure S4. The acetonitrile washes did not improve the activity of CFPS following DMSO exposure, suggesting that DMSO likely decreases activity by disabling some component of the CFPS during the exposure rather than by residual solvent impacting the reaction.

### Supplementary Materials and Methods

*PANOx-SP lysate preparation*

*E. coli* BL21 DE3 star bacteria were pre-cultured from frozen stock in 100 mL 2xYPTG media in a 500 mL shake flask at 37°C, 250 rpm for 16 hours. 5 mL of the pre-culture was used to seed 1 L of 2xYPTG media in a 2 L baffled shake flask that was shaken at 37°C, 250 rpm for approximately 3.25 hours, or until optical density at 600 nm reaches 3.0. Cells were then pelleted by centrifugation at 5000xg for 15 minutes and washed four times with S30A buffer (10 mM Tris acetate pH 8.2, 14 mM magnesium acetate, 60 mM potassium acetate, 2 mM DTT). The wet weight of the pellet was then measured before the pellet was flash frozen in liquid nitrogen and stored at -80°C until further processing. For lysis, cell pellets were re-suspended in cold S30A buffer at 1 mL buffer per 1 g wet cell mass, then lysed by sonication on ice in 1.5 mL aliquots (Qsonica Q500, 20% amplitude, 40 seconds on, 59 seconds off, 540 J total energy). 4.5 µL 1 M DTT is added to each tube after sonication. Lysates were then centrifuged at 12,000xg for 10 minutes at 4°C and the supernatant was collected, aliquoted, flash frozen in liquid nitrogen, and stored at -80°C until reaction assembly.

*3-PGA lysate preparation*

*E. coli* BL21 Rosetta2 bacteria were pre-cultured 7 hours at 37°C, 250 rpm from a colony in 3 mL 2xYPT media in a 14 mL culture tube. 30 µL of the pre-culture were then transferred to inoculate a “midi-culture” in 60 mL 2xYPT media in a 500 mL Erlenmeyer flask and was incubated 7 hours at 37°C, 250 rpm. Then, four 750 mL 2xYPT final cultures in 2.5 L baffled flasks are each inoculated with 7.5 mL of the midi-culture. Flasks are incubated with shaking at 37°C, 250 rpm for three hours or when optical density at 600 nm reaches 1.5-1.6. Cells were then pelleted at 5000xg for 10 minutes and washed four times with S30A* buffer (50 mM Tris pH 8.2, 14 mM magnesium glutamate, 60 mM potassium glutamate, 2 mM DTT). Note this S30A* buffer differs from the PANOx-SP S30A buffer by using magnesium and potassium salts with glutamate counter ion. The wet weight of the cell mass is measured, then diluted by a factor of 2.05mL per gram with S30A* buffer. A M-110P microfluidizer set to 16,000 psi is used to lyse the cells (Microfluidics, Westwood, MA). The lysate is centrifuged at 18000 rpm, 20 min, 4°C. The supernatant is pooled, then incubated 90 minutes at 37°C in a run-off reaction. Then, the lysate is centrifuged again at 20,000 rpm, 10 min, 2°C. The lysate supernatant is dialyzed for 1 hour at 4°C against 950 mL S30B buffer (14 mM magnesium glutamate, 150 mM potassium glutamate, 2M Tris pH 8.2, 1 mM DTT) before a last centrifugation at 20,000 rpm, 10 min, 2°C. The treated lysate supernatant is aliquoted, flash frozen in liquid nitrogen, and stored at -80°C until reaction assembly.

*CFPS Reagent preparation*

For PANOx-SP, a pre-mix solution was assembled containing magnesium glutamate, potassium glutamate, ammonium glutamate, ATP, GTP, CTP, UTP, folinic acid, tRNAs, each of the 20 canonical amino acids, phosphoenolpyruvate (PEP), nicotinamide adenine dinucleotide (NAD), coenzyme A (CoA), spermidine, putrescine, oxalic acid, and HEPES buffer (pH 7.4). Concentrations of each component in the final reaction are listed in Table S4. This premix stock solution was aliquoted, flash frozen in liquid nitrogen, and stored at -80°C until use in CFPS reaction assembly. In addition, leftover stock solutions of each component were flash frozen alone and stored at -80°C for use in the CFPS component screen experiment. Three PANOx-SP ingredients not included in the premix are stored separately at -20°C until reaction assembly: purified T7 RNA polymerase (RNAP), murine RNase inhibitor (New England Biolabs), and purified plasmid DNA.

The reagents added to 3-PGA CFPS reaction are somewhat different than PANOx-SP. The pre-mix solution for 3-PGA contains ATP, GTP, CTP, UTP, folinic acid, tRNAs, NAD, CoA, Spermidine, HEPES buffer (pH 8), cyclic adenosine monophosphate (cAMP), 3-phosphoglyceric acid (3-PGA), and dithiothreitol (DTT). Concentrations of each component in the final reaction are listed in Table S4. This premix is also aliquoted, flash frozen in liquid nitrogen, and stored at -80°C. A stock solution of all 20 amino acids are aliquoted and stored separately at -80°C. Other component stock solutions are stored at -20°C: potassium glutamate, magnesium glutamate, polyethylene glycol (PEG), and T7 RNAP. A maltodextrin solution is prepared fresh before reaction assembly.

*Lyophilization of CFPS reactions*

To lyophilize CFPS reactions in a microplate, the 96 well plate is dipped in liquid nitrogen and transferred to a shelf-type lyophilizer (SP Scientific, VirTis Wizard 2.0) with initial temperature set to -40°C. When the lyophilizer chamber is sealed, the vacuum is initiated to reach approximately 200-300 mTorr. Then, primary drying is initiated with a shelf temperature of -20°C and maintained for four hours. The Secondary drying step changes the shelf temperature to 15°C, and is maintained overnight or approximately 17 hours. Microplate samples removed from the lyophilizer were immediately treated with solvent as indicated and rehydrated to monitor protein synthesis activity.

*Acetonitrile wash after DMSO exposure*

Acetonitrile washes after DMSO exposure were tested for improved solvent removal. After DMSO exposure as described above, the insoluble reaction material is washed twice with 100 µL washes of acetonitrile, which are removed by aspiration followed by evaporation under a vacuum for 20 minutes. Reactions were rehydrated as described.

*Preparation, acetone treatment, and analysis of larger scale CFPS powder*

Larger 250 µL scale PANOx-SP CFPS reactions were assembled on ice in 1.5-mL Eppendorf tubes. For larger scale reactions, we pre-expressed T7 RNA polymerase in the E. coli culture used to make cell extract rather than adding T7 RNAP purified separately. Also, we omitted the RNAse inhibitor ingredient. These cost-saving measures did not majorly impact productivity in solvent tolerance experiments. The tubes were flash frozen in liquid nitrogen and transferred to a FreeZone 4.5 Liter benchtop manifold lyophilizer set to 0.08 mBar with internal refrigeration at -105°C. Samples were lyophilized for 48 hrs. The dry weight of the sample and tube were measured, and the weight of the tube alone was subtracted to find the dry weight of the CFPS mixture. The lyophilized CFPS cake was easily ground to a powder using a pipet tip. This powder was weighed into aliquots for TGA analysis. To compare acetone tolerance to fresh reactions, a 250 µL CFPS mixture was lyophilized and milled as above, submerged in 200 µL of acetone, and rehydrated in 250 µL of nuclease-free water. 5 uL aliquots of the rehydrated solution were distributed into clear, v-bottom 96 well plates to track GFP fluorescence using a SpectraMax M5 Microplate Reader. Reactions were carried out over the course of 4 hours at 30°C. Fluorescence signal was monitored with an ex/em of 490/515 nm with gain set to ‘low’. GFP signal in relative fluorescence units (RFU) was converted to µM GFP using fluorescence measurements of a purified GFP standard at the same settings on the microplate reader.

*PANOx-SP CFPS component screen*

CFPS components were screened to identify which components of the lysate-based PANOx-SP CFPS method could contribute to protect lyophilized lysate during acetone exposure. The component stock solutions tested are DNA, T7 RNAP, RNAse inhibitor, 10xSS, HEPES, 15xMM, 20AA, PEP, NAD, CoA, oxalic acid, putrescine, and spermidine. 10xSS refers to a solution of magnesium, potassium, and ammonium glutamate. “15xMM” is a solution of ATP, GTP, UTP, CTP, folinic acid, and tRNA. “20AA” is a solution containing all 20 canonical amino acids. Control reactions were labeled “None”, referring to PANOx-SP cell extract without any additional ingredients, and “Complete,” a complete PANOx-SP CFPS reaction mixture. Each component stock solution was stored frozen separately until reaction assembly.

To assemble reactions, each component listed above was first combined with lysate and water to reach a volume of 15 µL per well in the microplate.  The plate was flash frozen in liquid nitrogen and lyophilized as described above. 100 µL acetone was added to solvent wells for one hour while control wells were left untreated. A polypropylene mat was used as before to prevent solvent evaporation. Following acetone treatment, the majority of the solvent was removed by aspiration with care not to disturb the insoluble pellet. The remaining solvent was allowed to evaporate at room temperature in a fume hood (without application of a vacuum). Then, all reactions apart from the "complete" reaction controls were rehydrated with a mixture of water and all components of a complete CFPS reaction excluding the lysate (pre-mix, T7 RNAP, RNase inhibitor, and DNA stock solutions). "complete" control wells were rehydrated with water. This experimental setup results in a final concentration two times that of a normal reaction for the component being tested. For example, in the wells testing the DNA component, DNA is at a 1x6.4 nM concentration prior to lyophilization, but at a 2x6.4 nM concentration after rehydration with water and CFPS reagents. Acetone treatment results are normalized to the no-solvent control to isolate the effect of each added component on acetone sensitivity of the lysate.

### Supplementary Figures

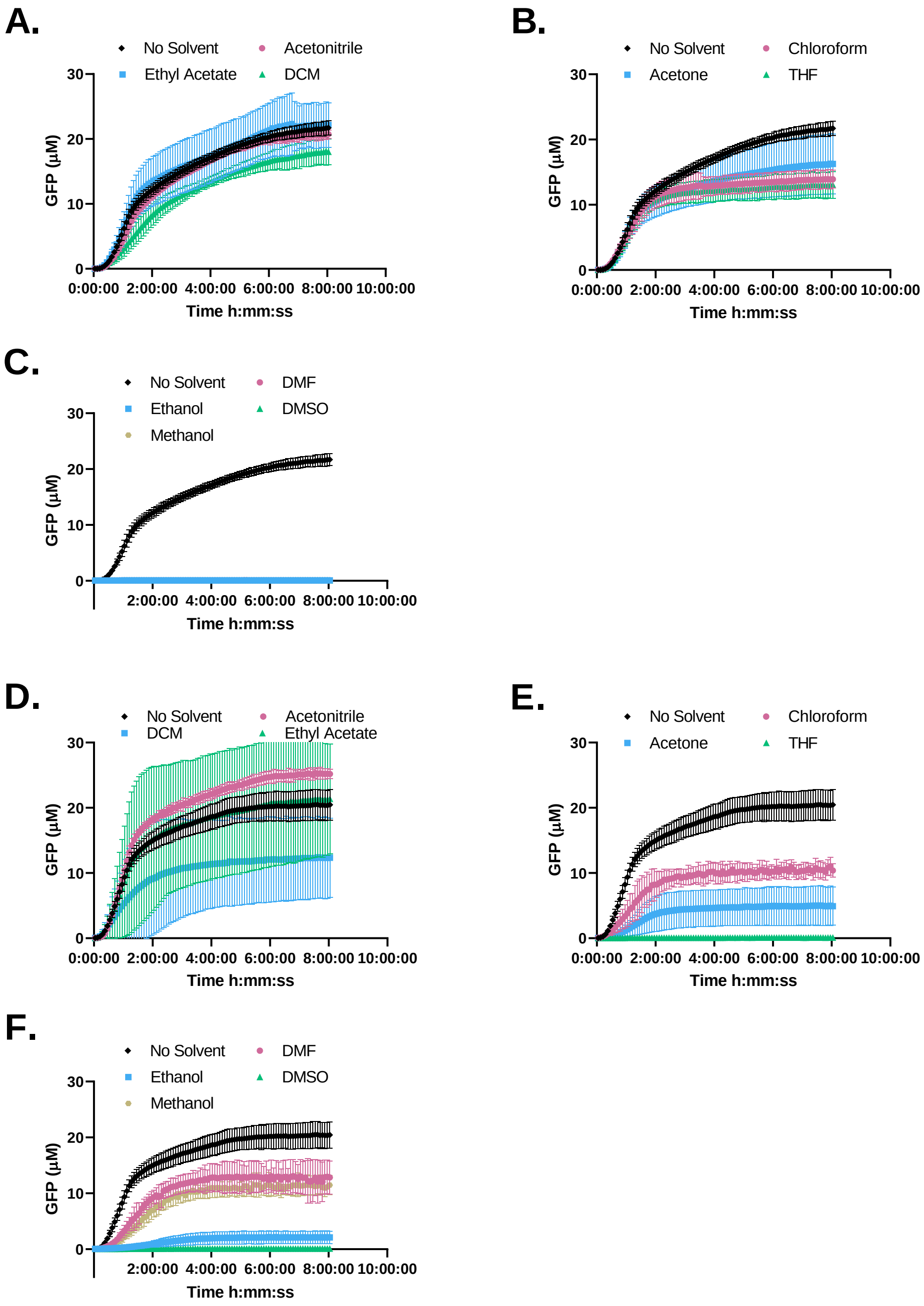

**Figure S1** Kinetics plots of solvent screen with endpoints represented in Figure 2 of the main text. Complete reactions exposed to solvent (a-c), cell lysate only exposed to solvent (d-f).

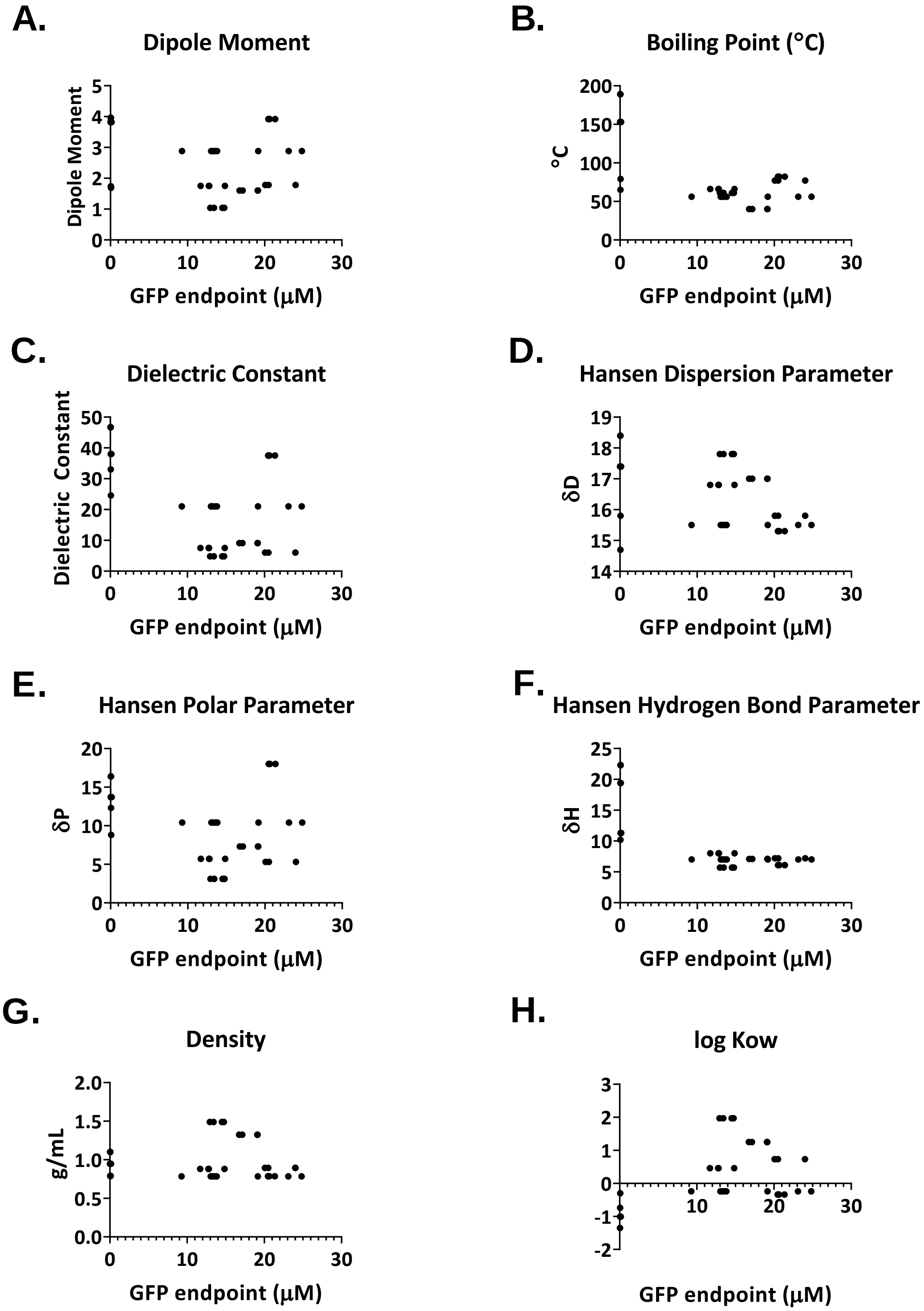

**Figure S2** Correlation plots comparing endpoint GFP productivity to the physical property parameters of the organic solvents used. Lyophilized, complete CFPS reactions were exposed to solvent, then rehydrated with water.

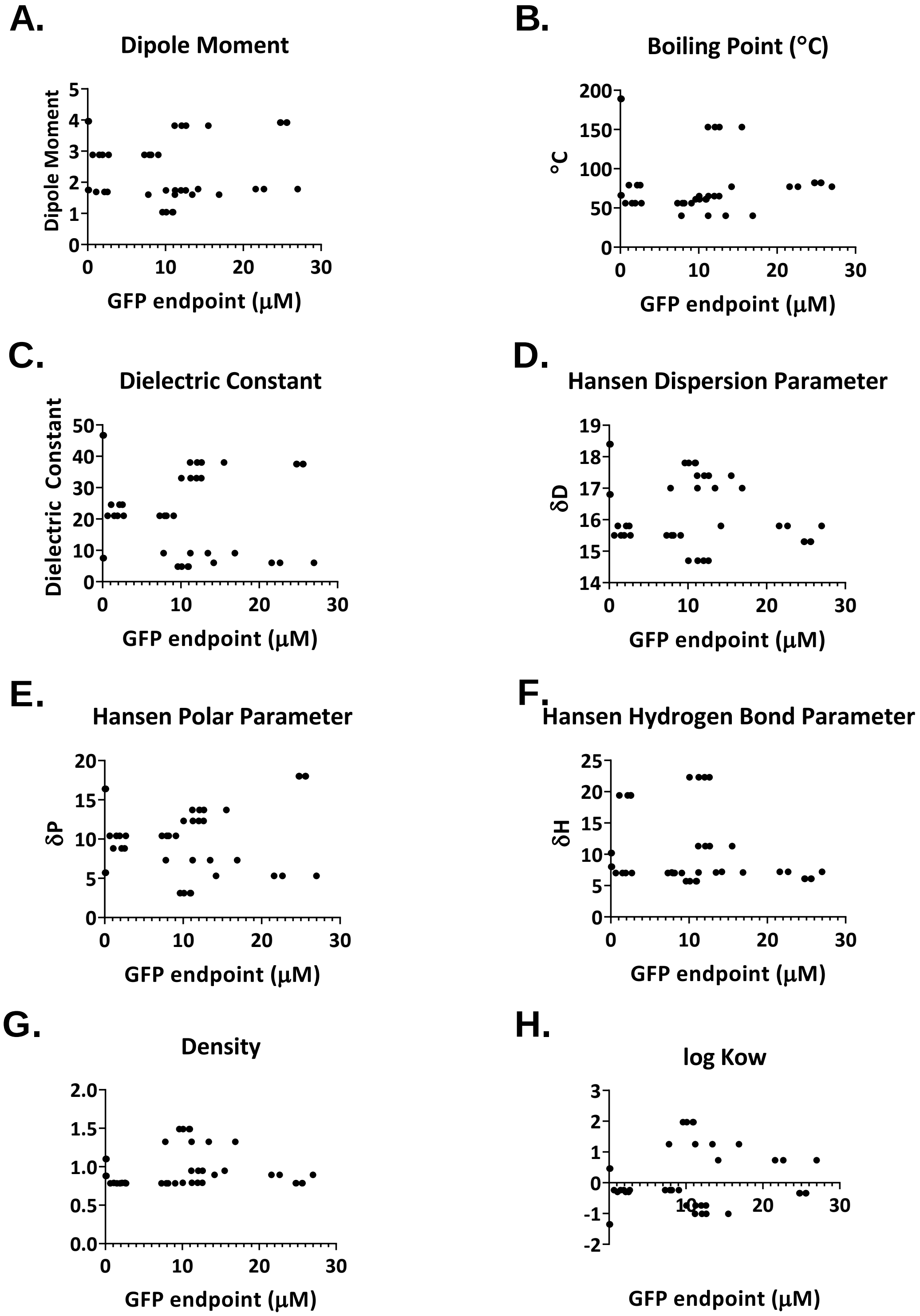

**Figure S3** Correlation plots comparing endpoint GFP productivity of solvent-exposed lysate to the physical property parameters of the organic solvents used. Lyophilized lysate was exposed to solvent alone, then complemented with the other ingredients of a complete CFPS reaction upon rehydration.

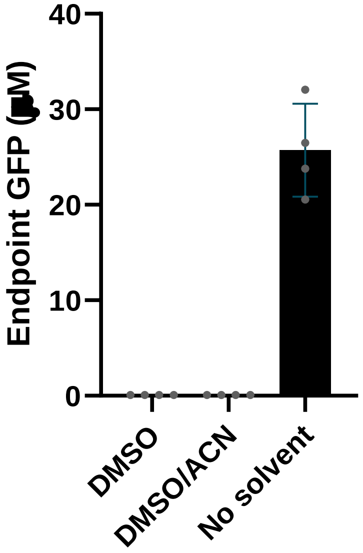

**Figure S4** Washing with acetonitrile to remove DMSO. Each bar across the x-axis is labeled with the solvent treatment. “DMSO/ACN” refers to samples treated first with DMSO, then washed with acetonitrile.

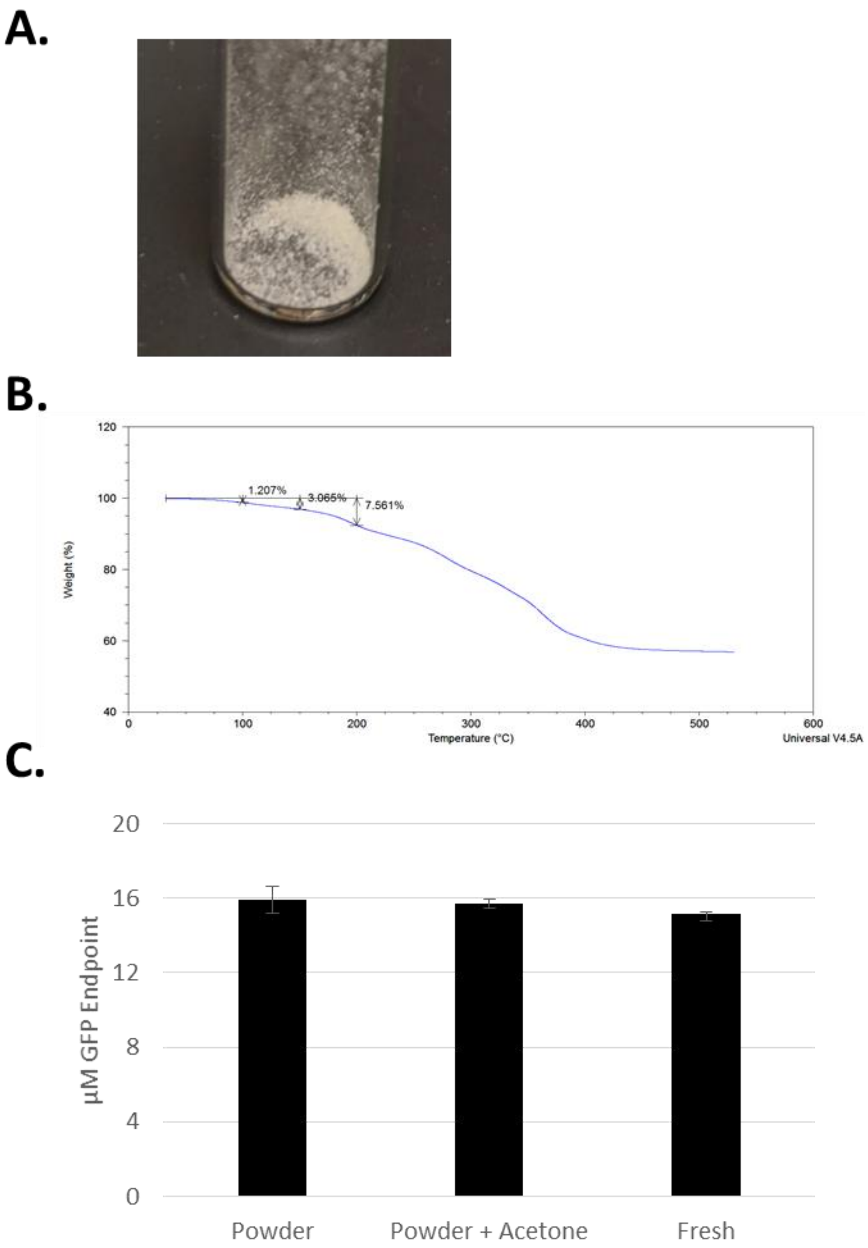

**Figure S5** CFPS powder after 48 hr lyophilization (a). TGA analysis of the powder (b) where water is the first volatile component and makes up 1.2-3% of the initial weight. Endpoint GFP concentrations after rehydration of CFPS powder and incubation for 8 hours, with and without acetone treatment (c). Non-lyophilized control reaction is labeled "fresh."

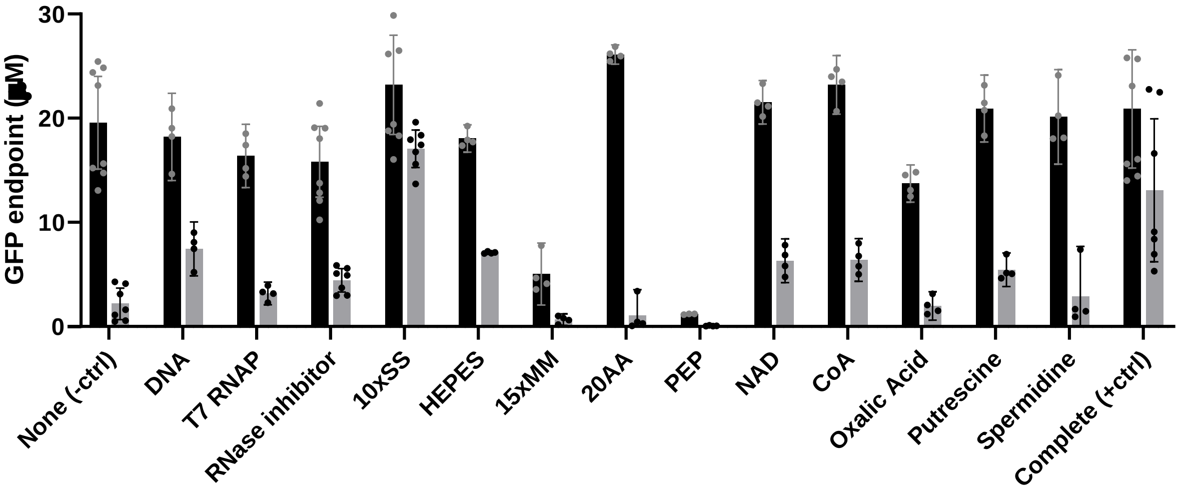

**Figure S6** Non-normalized PANOx-SP CFPS component screen for acetone tolerance. Black bars are results from a control without solvent exposure. Grey bars refer to a 1 hr acetone exposure. Acetone is evaporated under ambient conditions. Each set of bars is labeled with the CFPS ingredient used to supplement the lysate sample at drying. The negative control labeled “None” is lysate without any supplemented additive. The positive control labeled “Complete” includes all CFPS ingredients.

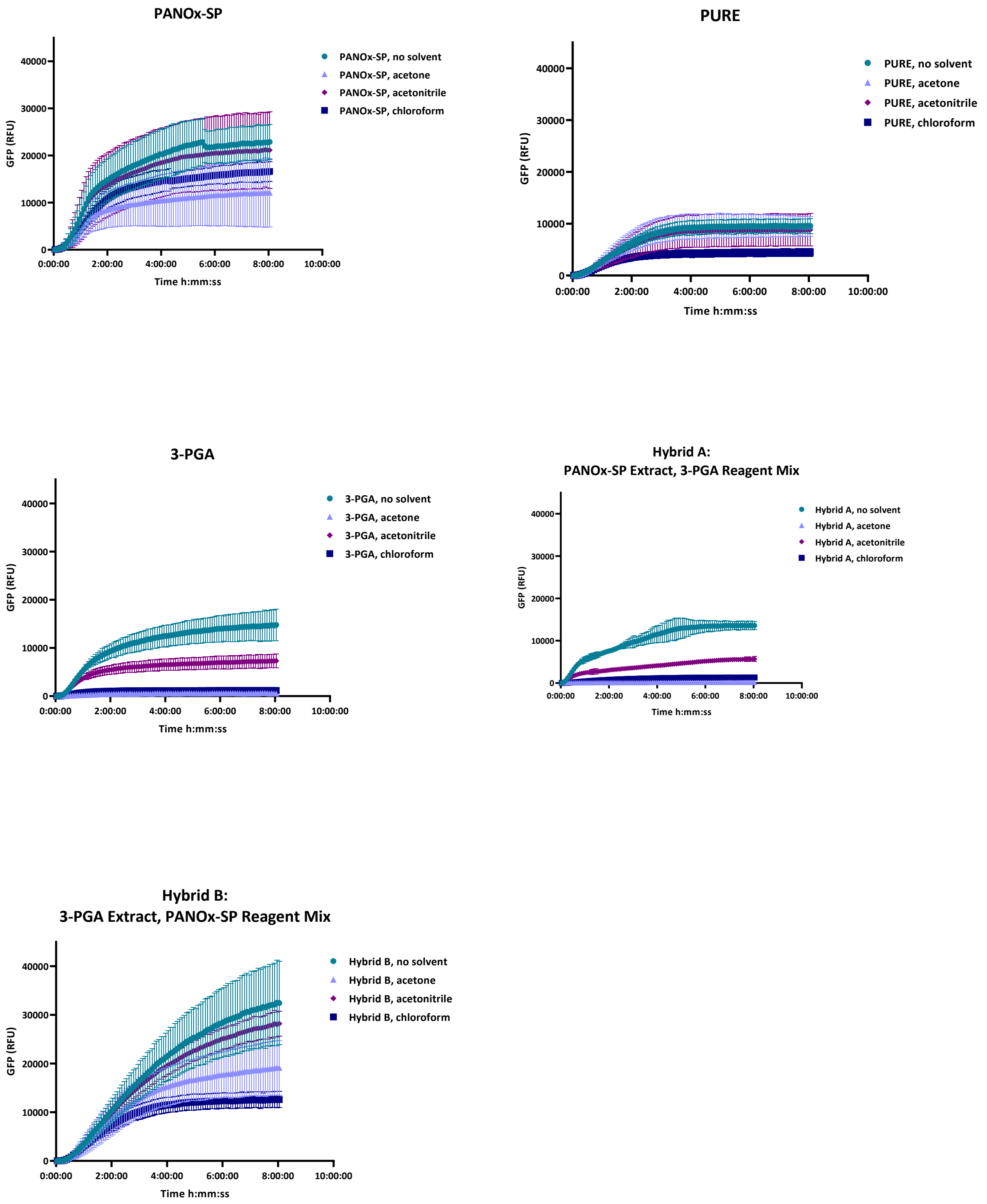

**Figure S7** Kinetics plots for different CFPS recipes exposed to four solvent conditions. Plots correspond to Figure 4 endpoint data in the main text.

### Supplementary Tables

**Table S1** Solvent properties and source information

| Solvent | Vendor | Catalog No. | Solvent Group | Dipole Moment | Boiling Point (°C) | Dielectric Constant | Hansen Dispersion Parameter | Hansen Polar Parameter | Hansen Hydrogen Bond Parameter | Density | Log(Kow) |
|---|---|---|---|---|---|---|---|---|---|---|---|
| **Acetone** | Millipore -Sigma | 270725 | Polar aprotic | 2.88 | 56 | 21 | 15.5 | 10.4 | 7.0 | 0.7845 | -0.240 |
| **Aceto-nitrile** | Millipore -Sigma | 360457 | Polar aprotic | 3.92 | 82 | 37.5 | 15.3 | 18 | 6.1 | 0.7860 | -0.340 |
| **Ethyl Acetate** | Acros | 42368-0040 | Polar aprotic | 1.78 | 77 | 6.02 | 15.8 | 5.3 | 7.2 | 0.8940 | 0.730 |
| **Dimethyl-formamide** | Millipore -Sigma | 186317 | Polar aprotic | 3.82 | 153 | 38 | 17.4 | 13.7 | 11.3 | 0.9480 | -1.010 |
| **Dichloro-methane** | Millipore -Sigma | D65100 -4L | Non-polar | 1.60 | 40 | 9.1 | 17.0 | 7.3 | 7.1 | 1.3250 | 1.250 |
| **Tetrahydro -furan** | Millipore -Sigma | 34865-100mL | Polar aprotic | 1.75 | 66 | 7.5 | 16.8 | 5.7 | 8 | 0.8800 | 0.460 |
| **Ethanol** | Millipore -Sigma | 493511 -4L | Polar protic | 1.69 | 79 | 24.55 | 15.8 | 8.8 | 19.4 | 0.7893 | -0.300 |
| **Methanol** | Fisher | A411-4 | Polar protic | 1.74 | 65 | 33 | 14.7 | 12.3 | 22.3 | 0.7910 | -0.740 |
| **Chloroform** | Fisher | C6060-1 | Non-polar | 1.04 | 61 | 4.81 | 17.8 | 3.1 | 5.7 | 1.4900 | 1.970 |
| **DMSO** | Sigma | D2650 | Polar aprotic | 3.96 | 189 | 46.7 | 18.4 | 16.4 | 10.2 | 1.1000 | -1.350 |

**Table S2** Correlation statistics for productivity of solvent-exposed CFPS reactions

| **GFP endpoint (µM) Complete reaction exposure** | **vs. Dipole Moment** | **vs. Boiling Point** | **vs. Dielectric Constant** | **vs. Hansen Dispersion** | **vs. Hansen Polar** | **vs. Hansen Hydrogen Bond** | **vs. Density** | **vs. log(Kow)** |
|---|---|---|---|---|---|---|---|---|
| **Pearson r** | | | | | | | | |
| r | -0.1556 | -0.5766 | -0.5830 | -0.2311 | -0.3196 | -0.7478 | 0.08675 | 0.5907 |
| 95% confidence interval | -0.4325 to 0.1481 | -0.7458 to -0.3375 | -0.7501 to -0.3460 | -0.4941 to 0.07063 | -0.5631 to -0.02514 | -0.8549 to -0.5796 | -0.2157 to 0.3740 | 0.3653 to 0.7552 |
| R squared | 0.02423 | 0.3325 | 0.3399 | 0.05341 | 0.1022 | 0.5592 | 0.007526 | 0.3489 |
| **P value** | | | | | | | | |
| P (two-tailed) | 0.3130 | <0.0001 | <0.0001 | 0.1312 | 0.0344 | <0.0001 | 0.5755 | <0.0001 |
| P value summary | ns | **** | **** | ns | * | **** | ns | **** |
| Significant? (alpha = 0.05) | No | Yes | Yes | No | Yes | Yes | No | Yes |
| **Number of XY Pairs** | 44 | 44 | 44 | 44 | 44 | 44 | 44 | 44 |

**Table S3** Correlation statistics for productivity of solvent-exposed *E. coli* lysate

| **GFP endpoint (µM) Complete reaction exposure** | **vs. Dipole Moment** | **vs. Boiling Point** | **vs. Dielectric Constant** | **vs. Hansen Dispersion** | **vs. Hansen Polar** | **vs. Hansen Hydrogen Bond** | **vs. Density** | **vs. log(Kow)** |
|---|---|---|---|---|---|---|---|---|
| **Pearson r** | | | | | | | | |
| r | 0.09429 | -0.1509 | -0.02309 | -0.3073 | 0.1437 | -0.2137 | -0.002955 | 0.1932 |
| 95% confidence interval | -0.2084 to 0.3805 | -0.4286 to 0.1528 | -0.3178 to 0.2757 | -0.5537 to -0.01151 | -0.1600 to 0.4226 | -0.4801 to 0.08882 | -0.2996 to 0.2942 | -0.1100 to 0.4635 |
| R squared | 0.008891 | 0.02277 | 0.0005334 | 0.09446 | 0.02065 | 0.04566 | 8.730e-006 | 0.03732 |
| **P value** | | | | | | | | |
| P (two-tailed) | 0.5426 | 0.3282 | 0.8817 | 0.0424 | 0.3521 | 0.1637 | 0.9848 | 0.2089 |
| P value summary | ns | ns | ns | * | ns | ns | ns | ns |
| Significant? (alpha = 0.05) | No | No | No | Yes | No | No | No | No |
| **Number of XY Pairs** | 44 | 44 | 44 | 44 | 44 | 44 | 44 | 44 |

**Table S4** Final concentrations of components in three different CFPS buffers

| Component | PANOx-SP | 3-PGA | PURE |
|---|---|---|---|
| Magnesium glutamate | 12 mM | 10 mM + 4.62 mM† | - |
| Potassium glutamate | 130 mM | 30 mM + 49.5 mM† | 100 mM |
| Ammonium glutamate | 10 mM | - | - |
| Magnesium acetate | 3.7 mM* | - | 13 mM |
| Potassium acetate | 16.02 mM* | - | - |
| Tris acetate | 2.67 mM (pH 8.2)* | 1.65 mM (pH 8.2)† | - |
| HEPES | 57 mM (pH 7.4) | 50 mM (pH 8) | 50 mM (pH 7.6) |
| ATP | 1.2 mM | 1.5 mM | 2 mM |
| GTP | 0.85 mM | 1.5 mM | 2 mM |
| CTP | 0.85 mM | 0.9 mM | 1 mM |
| UTP | 0.85 mM | 0.9 mM | 1 mM |
| Folinic acid | 0.072 mM | 0.068 mM | 0.02 mM |
| tRNA | 170.6 µg/mL | 200 µg/mL | 2.8 $A_{260}$ units in 50 µL = 3.5 mg/mL |
| Alanine | 2 mM | 3 mM | 0.3 mM |
| Arginine | 2 mM | 3 mM | 0.3 mM |
| Histidine | 2 mM | 3 mM | 0.3 mM |
| Lysine (monoHCl) | 2 mM | 3 mM | 0.3 mM |
| Aspartic acid | 2 mM | 3 mM | 0.3 mM |
| Glutamic acid | 2 mM | 3 mM | 0.3 mM |
| Isoleucine | 2 mM | 3 mM | 0.3 mM |
| Leucine | 2 mM | 3 mM | 0.3 mM |
| Methionine | 2 mM | 3 mM | 0.3 mM |
| Phenylalanine | 2 mM | 3 mM | 0.3 mM |
| Tryptophan | 2 mM | 3 mM | 0.3 mM |
| Tyrosine | 2 mM | 3 mM | 0.3 mM |
| Valine | 2 mM | 3 mM | 0.3 mM |
| Serine | 2 mM | 3 mM | 0.3 mM |
| Threonine | 2 mM | 3 mM | 0.3 mM |
| Asparagine | 2 mM | 3 mM | 0.3 mM |
| Glutamine | 2 mM | 3 mM | 0.3 mM |
| Cysteine | 2 mM | 3 mM | 0.3 mM |
| Glycine | 2 mM | 3 mM | 0.3 mM |
| Proline | 2 mM | 3 mM | 0.3 mM |
| PEP | 33 mM | - | - |

| | | | |
|---|---|---|---|
| NAD | 0.33 mM | 0.33 mM | - |
| CoA | 0.27 mM | 0.26 mM | - |
| Spermidine | 1.5 mM | 1 mM | 2 mM |
| Putrescine | 1 mM | - | - |
| Oxalic acid | 4 mM | - | - |
| T7 RNA polymerase | 100 µg/mL | 100 µg/mL | 10 µg/mL |
| Plasmid DNA | 6.4 nM | 6.4 nM | 6.4 nM |
| RNase Inhibitor | 0.8 U/µL | - | - |
| Cell extract | 26.7% v/v* | 33% v/v† | - |
| Maltodextrin | - | 30 mM | - |
| PEG | - | 1.5 % w/v | - |
| cAMP | - | 0.75 mM | - |
| 3-PGA | - | 30 mM | - |
| DTT | 0.8 mM* | 1 mM+0.33mM† | 1 mM |
| Creatine Phosphate | - | - | 20 mM |

* Items in Style I recipe added as part of the cell extract solution

† Items in Style II recipe added as part of the cell extract solution
